# Quality assessment for different haplotyping methods and GWAS sensitivity to phasing errors

**DOI:** 10.1101/015669

**Authors:** Giovanni Busonera, Marco Cogoni, Gianluigi Zanetti

## Abstract

In this report we present a multimarker association tool (Flash) based on a novel algorithm to generate haplotypes from raw genotype data. It belongs to the entropy minimization class of methods [4, 7] and is composed of a two stage deterministic - heuristic part and of a optional stochastic optimization. This algorithm is able to scale up well to handle huge datasets with faster performance than the competing technologies such as BEAGLE[5] and MACH[10] while maintaining a comparable accuracy. A quality assessment of the results is carried out by comparing the switch error. Finally, the haplotypes are used to perform a haplotype-based Genome-wide Association Study (GWAS). The association results are compared with a multimarker and a single SNP association test performed with Plink [12]. Our experiments confirm that the multimarker association test can be more powerful than the single SNP one as stated in the literature. Moreover, Flash and Plink show similar results for the multimarker association test but Flash speeds up the computation time of about an order of magnitude using 5 SNP size haplotypes.

## 1. Introduction

Genome-wide association studies (GWAS) are used to identify common genetic factors that influence health and disease. Basically GWAS are performed at the single nucleotide level. In Ballard et al.[3] is stated that the joint use of information from multiple markers may be more effective to reveal association between a genomic region and a trait than single marker analysis. Since a multimarker GWAS requires the reconstruction of the phase from genotyped data, which is usually a time expensive task, a fast haplotyping algorithm is crucial.

In this report we present a scalable, fast and reasonably accurate haplotype-based association tool that reconstructs haplotypes used to perform a multi-marker chi-square association test.

Unlike most phasing tools, based on a statistical approach (i.e Hidden Markov Model), our method is composed of a two stage deterministic-heuristic part and of a optional stochastic optimization.

The work is organized as follows. In Section 2 the new algorithm is sketched. Section 2.1 is devoted to the description of the deterministic initial part of the method and to show that it is able to efficiently compute haplotypes up to 6 SNP. In Section 2.2 a stochastic approach to phasing is described that turns out to be useful when windows are larger than 6 SNPs. *A possible extension of the algorithm to work with a dynamic window size by computing the linkage disequilibrium (LD) on the fly is planned*.

Numerical tests to check for quantitative differences among different haplotyping techniques are in Section 4 where a thorough comparison of phasing results is presented.

The final section focuses on GWAS performed both on single SNPs and haplotype-based. The main goal is to show that association results are not influenced by small differences among inferred haplotypes.

## 2. Description of the FLASH algorithm

The Flash algorithm is based on a minimization approach to phase a given set of genotypes. It exploits the correlation among individual SNP data to find the minimum set of haplotypes that describe them. At the same time the algorithm tries to minimize the entropy of the solution.

*At the moment, the algorithm phases consecutive segments of the chromosome by means of a fixed size sliding window*. Depending on the windows size a totally deterministic approach, which explores the whole solution space to find the optimal one, is not feasible. To cope with this problem two algorithms were developed and implemented: a heuristic based (baseline algorithm) and a stochastic based (Simulated annealing). The former one is more accurate but can reach window size of about 6 SNP on a common workstation. The latter can work with larger window sizes with a small accuracy loss.

### 2.1. Baseline Algorithm

For the generic segment *seg* some data structures are defined:

1. The Individual Genotypes list *G_seg_* holds the genotypes related to each individual (ID).
2. The individual compatible diplotypes *D_seg_* map holds a list for each ID with all possible phased diplotypes (the haplotype couple).
3. The *M_seg_* diplotype matrix is built to store at the *i, j* position the ID solved by the *i, j* diplotype.
4. The haplotypes occurrence table *H_seg_* stores the occurrences of the haplotypes that are in *D_seg_*.
5. The solution set *S_seg_* contains the solving haplotypes.
6. The unsolved individuals *U_seg_* set holds the IDs temporarily unsolved.
7. The legal haplotype set (if needed) *L_seg_* holds the haplotypes used to perform a heuristic search.
8. Extended haplotypes set *E_seg_* is used to extend *S_seg_* with a combination of haplotypes taken from *L_seg_*.

For each segment the algorithm performs the following steps:

1. **Computation of the** *D_seg_* **map:** All genotypes in *G_seg_* are analyzed and a list of compatible diplotypes is built. IDs with the same genotype in *G_seg_* are considered indistinguishable and will be solved with the same diplotype. The diplotype matrix *M_seg_* is also created.
2. *U_seg_* **initialization:** The *U_seg_* list is filled with all the IDs.
3. **Haplotype occurrence computation:** for each haplotype in the *D_seg_* map the occurrence is computed and stored in *H_seg_*.
4. **Computation of** *S_seg_* **with mandatory haplotypes:** IDs with only one occurrence in *M_seg_* can be solved only with one or two mandatory haplotypes. These haplotypes are used to create *S_seg_*. IDs solved are removed from *U_seg_*. If no IDs have mandatory haplotypes *S_seg_* ≡ Ø.
5. **Iteration to solve unsolved people using haplotypes added to** *S_seg_:* If *S_seg_* ≠ Øand *U_seg_* ≠ Ø the algorithm tries to solve remaining IDs using haplotypes in *S_seg_*. When an ID is solved it is removed from *U_seg_*. If *U_seg_* ≡ Ø the computation is completed and all the IDs are then phased using haplotypes in *S_seg_*.
6. **An heuristic phase search in the configuration space to find the complete solution (if needed):** If *U_seg_* ≠ Ø after the previous steps means *S_seg_* set has to be augmented with other haplotypes to solve the global problem. To find them the following heuristic algorithm is used:
  a. Consider all the*h_i_* haplotypes taken from diplotype compatible solutions of IDs still in *U_seg_*.
  b. Create the legal set *L_seg_* with *h_i_* : *h_i_* ∉ *S_seg_*. Sort *h_i_* in *L_seg_* with respect to their occurrence stored in *H_seg_*. Let *M* be the cardinality of *L_seg_*.
  c. For *n* ∈ [1 : *M*]:
    i. For each combination C_*in*_ of *n* haplotypes in *L_seg_*:
      A. Create the extended haplotype set *E_seg_ = S_seg_* ∪ *C_in_*.
      B. If *E_seg_* haplotypes can describe all the IDs in <u>*U_seg_*</u> the problem is considered solved. The algorithm is completed.
      C. If *E_seg_* haplotypes is not able to describe the IDs in <u>*U_seg_*</u> keep on iterating among combinations.

A brief analysis of the algorithm provides an upper bound of the total iterations needed to phase a single segment of *W* SNPs. Even though the different possible genotypes are 3*^W^*, the maximum number of haplotypes that can phase the segment is 2*^W^*. Now, suppose that there is no genotype with mandatory symbols. The *S_seg_* ≡ Ø and the *L_seg_* set contains all compatible haplotypes. If the global solution is *S_seg_* ≡ *L_seg_*, the number of iterations needed to compute it, is

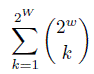

This worst case is shown in the table 1 for different values of *W:*

**Table 1.**
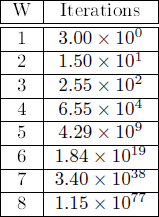
Number of iterations of the worst case

During the development of the algorithm we observed that in most cases a large number of individuals (over 90%) are solved in a relatively small number of iterations while reaching the totality of the solution often takes a lot of time. This is probably due to the fact that a small set of individuals have to be solved with low frequency haplotypes making the algorithm iterate among the less likely solutions. To give flexibility to the algorithm a feature to exit from the heuristic step when a given ratio of solved individuals or computation effort is reached, has been implemented. The computation effort is defined as the ratio between the current iteration and the theoretical maximum iterations as given by Table 1.

As an example of the baseline algorithm, consider a single segment with the 3-SNPs genotypes specified in table 2.

**Table 2.** Compatible diplotype table for the example case. The first column refers to the individual ID, the second one to the related genotype. The third column shows the possible phasing solutions for the ID.

| ID | Genotype | Compatible diplotypes |
| --- | --- | --- |
| 0 | 010,010 | 010,010 |
| 1 | 100,100 | 100,100 |
| 2 | 001,010 | 001,010 - 000,011 |
| 3 | 001,001 | 001,001 |
| 4 | 100,101 | 100,101 |
| 5 | 011,101 | 011,101 - 001,111 |

Their alleles are coded using digits 1 and 0. The third column shows the possible diplotypes that “solve” the phasing problem for that ID. The solutions differing for just the initial phase are considered equal (for instance 001,010 is equal to 010, 001).

The corresponding diplotype matrix is shown in table 3:

**Table 3.** Diplotype matrix for the example case. The *i, j* cell holds the individual ID solved by the *i, j* diplotype.

|  | 000 | 001 | 010 | 011 | 100 | 101 | 110 | 111 |
| --- | --- | --- | --- | --- | --- | --- | --- | --- |
| 000 |  |  |  | 2 |  |  |  | 5 |
| 001 |  | 3 | 2 |  |  |  |  |  |
| 010 |  |  | 0 |  |  |  |  |  |
| 011 |  |  |  |  |  | 5 |  |  |
| 100 |  |  |  |  | 1 | 4 |  |  |
| 101 |  |  |  |  |  |  |  |  |
| 110 |  |  |  |  |  |  |  |  |
| 111 |  |  |  |  |  |  |  |  |

At first, the haplotype occurrences are computed (see table 4).

**Table 4.** Haplotype occurrence table for the example case. For each haplotype, the number of occurrences among all the possible compatible diplotypes shown in the third column of table is computed.

| Haplotype | Occ. |
| --- | --- |
| 000 | 1 |
| 001 | 4 |
| 010 | 3 |
| 011 | 2 |
| 100 | 3 |
| 101 | 2 |
| 110 | 0 |
| 111 | 1 |

Then the solution alphabet is populated with the mandatory haplotypes. So at the first step we have:

*S* = {001, 010,100,101}

*U* = {2, 5}

Since *U* ≠ Ø, the algorithm tries to solve the remaining IDs with the haplotypes in *S*.

At this point, only the ID 2 has a compatible diplotype (001,010) with both haplotypes contained in *S*, so it can be marked as solved. Unfortunately, the ID 5 has just one haplotype in the set *S* for each compatible diplotype and therefore cannot be solved. To find a haplotype that completes the solution for the ID 5, the heuristic step is then invoked.

The legal set *L* is built using the haplotypes taken from the set of compatible diplotypes of ID 5 not present in *S*. This set is then sorted with respect to the haplotype occurrences stored in the *H* table (i.e. *L* = {011,111}). At this point the *E* set is created using the first of the combinations of one haplotype (*C*_11_ = {011}) from the *L* set: *E* = *S* ∪ *C*_11_ = {001,010,100,101,011}. At this point at least one compatible diplotype for each ID has both haplotypes in the set *E*. The final solution is then *S* ≡ *E*.

It is worth to note that the algorithm stops once a solution is found. It is not an actual brute force approach since the other possible *E* sets built using all the L combinations (*S*∪ {111},*S*∪ {011, 111}) are not analyzed even though are all valid solutions. This heuristic approach, however, provides good results reducing computation time. Moreover, sorting the legal set *L* allows often to find the minimum entropy solution among the ones with the same number of haplotypes.

### 2.2. Simulated annealing (SA) optimization

The method described in the previous section is usually capable of quickly find the smallest set of symbols able to describe the whole population. When the window size is larger than *s* = 6 the space of possible symbol combinations grows to a computationally prohibitive size (see table 1). An exhaustive search of such a space of combinations is inherently inefficient and not very clever. This problem is reminiscent of the estimation of the average value of some physical quantity for systems described by statistical mechanics in some thermodynamic ensemble.

Trying to mimic the statistical mechanics approach, we drop the deterministic method for finding the best solution and exploit a stochastic strategy in which some functional is optimized. To solve this problem the quantity to be minimized is the entropy of the haplotype solution. Here the definition of the Shannon entropy is used (*H* = – ∑*_i_ p_i_* log_2_ *p_i_* where *p_i_* is the probability of the i-th haplotype and *S* can also be seen as the average value of the logarithm of *p*).

As described in Section 2.1, at the beginning of the haplotyping procedure, the set *S_seg_* is generated by selecting the strictly necessary haplotypes. These haplotypes can be seen as degrees of freedom that the problem loses at the very beginning. This is done in a completely deterministic fashion.

In the next stage of the process we are left with a subset of individuals *U_seg_* not solved yet by the haplotypes present in *S_seg_*. The goal is to find a complementary set *C_seg_* of extended haplotypes which can be built in a stochastic manner by starting with an educated guess among ID diplotypes (choosing them in a random fashion makes the convergence slower). This part of the process replaces the one described at point 6 of Section 2.1. The main difference relies essentially in how the configurations space is explored: no more iteration over the legal haplotype combinations, but generation of random moves trying to find the extremal minimal entropy:

1. When a “fully legal” solution compatible with the starting genotypes of all the individuals has been prepared, the optimization process can begin by computing its entropy *H*_0_ and setting an initial temperature *T*_0_;
2. The previous solution is modified in a way that it is still legal but one of the individuals chosen at random (and eventually those compatible with it) is described by a different haplotype. This random choice of the haplotypes can be uniform or can be biased depending on the relative occurrence of each haplotype;
3. The entropy of the proposed modified solution *H_mod_* is computed and:
  a. if it is lower than the entropy of the previous state (at the beginning *H*_0_) it becomes the new current solution;
  b. if it is higher, it is accepted with some relatively small probability when a random number *r* ∈ [0,1) is larger than exp(Δ*H*/*kT*). So it is possible to accept a solution with a larger *H* depending on the entropy change Δ*H* and on the temperature parameter. The temperature should slowly go to zero within a reasonable amount of iterations as if the system would undergo an annealing process.
4. If the trend in the entropy of the solutions meets some empirical scheme for reached stability, the solution is considered the final one. Instead the algorithm goes back to the point 2 to get a new modified solution.

### 2.3. Missing data management

So far it was assumed that genotypes were given without missing data. In real datasets this situation is very unlikely and a certain amount of missing data is always present. During the haplotyping process an imputation phase is mandatory to fill the nocall (NC) sites. The Flash baseline algorithm and the SA use two different approaches to impute data. The following subsections describe both situations.

#### 2.3.1. Baseline

Missing SNPs in a segment are managed using the following steps:

1. All IDs with one or more nocall loci are inserted in a list *NC_seg_*. They are not present in the *U_seg_* list discussed in 2.1.
2. After the baseline algorithm introduced in 2.1 has completed, the haplotypes in the global solution *S_seg_* are used to create compatible diplotypes for IDs in *NC_seg_* in order to phase them.
3. If after the previous step *NC_seg_* ≠ Ø, a process like the heuristic used in the baseline algorithm is performed. From each IDs in *NC_seg_* a list of compatible diplotype is created and the relative haplotypes are inserted in the *L_NC_seg__* set. This set is used in the same manner as the *L_seg_*set to build extended haplotype sets used to find a legal solution for all IDs in *NC_seg_*. The algorithm stops once *NC_seg_* is empty.

#### 2.3.2. Simulated annealing

Before the beginning of the SA optimization, every individual with a genotype segment containing one or more NC data is detected and it is treated as if it had multiple possible genotypes. These compatible genotypes are generated as a tree which has three branches every time a NC site is found. At this point the individual with a segment containing incomplete genotype data acts as a “degrees of freedom multiplier” with respect to the final global solution. Since the SA optimizer has the goal of entropy minimization, the description of the segments with NCs will be the one (among all the combinations of the tree just described) that guarantees the least global entropy. In other words, if among the branches of the tree there is some haplotype that has already been used as a diplotype solution for some other ID, it will be favoured against those symbols adding complexity to the global solution.

### 2.4. Missing features in the current Flash implementation

The current implementation of flash lacks of some features that will be added in the future versions:

- A stochastic management of genotypes dataset errors
- Imputation of the raw data using a high resolution reference panel
- Phasing with a LD driven dynamic size sliding window

These add-ons can improve phasing accuracy of Flash and also speed performance. A brief discussion on their implementation is made in the following subsections.

#### 2.4.1. Genotype data error management

Raw genotype data contains some loci marked as NC. Moreover every position in the genotype is labeled with a “level of confidence” parameter. Up to now the algorithm cannot exploit this information.

There are two main possible approaches:

- The NC code (see Sec. 2.3) could be generalized to include a continuous range of confidence *c* ∈ [0,1) (no calls have almost zero confidence by definition.) This solution impacts negatively on the whole efficiency of the process due to the large number of degrees of freedom added to the optimization process.
- Another way to exploit this information could be to post-process the result of the standard algorithm.

#### 2.4.2. Reference Panel

The imputation of data from a higher resolution reference panel helps the accuracy of the global phasing. To tackle this problem an algorithm was developed but at the moment it is not implemented yet in the Flash software package. A description of this algorithm can be summarized as follow:

1. From the reference panel take the haplotypes and put them in the *S_seg_* set. These haplotypes are considered as mandatory for our dataset.
2. Since the panel has an higher resolution, SNPs in a window of the dataset to be phased have to be considered not contiguous but with “holes” between them. These “holes” are marked as nocall loci.
3. The algorithm used for nocall data introduced in 2.3.1 is then used to impute missing data.

#### 2.4.3. Dynamic size sliding window

At the moment the haplotyping algorithm expects as input a raw genotyped data that has been cut into many fixed length *W* overlapping segments by means of a sliding window. Every couple of consecutive segments has exactly *W* – 1 SNPs in common. This procedure would work well if the raw data could be characterized by an homogeneous linkage disequilibrium value compatible with a fixed length window.

When very large datasets, possibly of different origin and with different SNP spacing distribution have to be haplotyped, having the ability to compute the local distribution of LD for every SNP location could be very useful. For instance, it would be possible to dynamically adapt the window size to the local value of LD.

## 3. Application of the method to haplotype based GWAS

A genome-wide association study (GWAS) is an examination of many common genetic variants in different individuals to see if any variant is associated with a phenotypical trait. GWAS typically focuses on associations between (sets of) SNPs and traits like major diseases.

In this kind of study there is usually a pool of genomes partly built from subjects showing the trait (cases) and others who do not (controls). If one allele is more frequent in people with the disease, the marker is said to be “associated” with the disease.

Most of the association studies are based on the calculation of the correlation between a single SNPs marker and the phenotype: this is the easiest and least computationally intensive approach. It has been demonstrated that an association study based not on single SNPs but on haplotypes can increase statistical power [11, 13, 2, 6]. This advantage means that the useful signal will be less buried in the noise in a Manhattan plot and usually a smaller p-value (along with a larger *χ*^2^) will be obtained from the analysis.

To exploit the improved statistical power of the multimarker procedure, for each phased segment we performed a haplotype based *χ*^2^ association test. The approach is similar to the one followed by Plink, but its phasing method is based on a estimation-maximization (EM). We perform both an OMNIBUS and a per haplotype association test. The details of this Flash application is shown in section 4.2.

## 4. Results

This section describes the tests performed to evaluate the haplotype inference performance and the ability to detect associations between haplotypes and phenotypes.

### 4.1. Haplotype estimation accuracy and computation performance

To evaluate the haplotype inference accuracy and computation performance, we compared the Flash and Beagle [5] results using both simulated and real data. The simulated data was generated using the MS [8] application whereas the real data were taken from the GENOME1000 project^1^. As a way to compare the accuracy of the phasing processes we employed the switch error defined as the proportion of successive pairs of heterozygote markers with incorrect phasing (in an individual) with respect to each other. As regards the computation performance, the time difference between the start of the application and the end of the data output is used.

The phasing tests are then performed using a set of overlapping SNPs windows (each window overlaps the adjacent ones for its size minus one) obtained slicing the input data accordingly.

The current implementation of Flash lacks the Simulated Annealing optimization (described in section 2.2) so haplotype inference obtains relevant speedups, with respect other similar applications like Beagle or Plink [12], when windows up to 5 SNPs wide are used.

Since Beagle lacks the feature to phase overlapping windows automatically, we performed two different Beagle runs for each dataset. The first one was performed on the whole dataset having the results sliced at the end of the computation in order to obtain the output in the same format as that produced by Flash. In the second run the dataset was sliced before the computation and a single instance of Beagle for each window was run. Since Beagle is based on a statistical approach (i.e Hidden Markov Model), slicing the dataset before the computation causes an information loss hindering the full Beagle performance, but it’s the only way allowing a parallel execution. In practice we found that, in the case of Beagle, the overhead of running a single instance of the application for each window, limits the advantages of running simultaneously on different nodes as shown in section 4.1.2.

#### 4.1.1. Simulated data

Each dataset, with different mutation and recombination rate (assigned with different MS parameter *t* and *r*), was created by simulating 100 different samples of 2000 haplotypes (made of 30 up to 300 SNPs). Results were then computed by averaging over 100 samples for each dataset. Since each sample contains relatively few SNPs, we did not perform the slicing before the computation, for the Beagle case, because we were mainly interested to evaluate the accuracy rather than the speed. By the way, performing real data tests, we discovered that running Beagle in parallel did not speedup the execution.

These tests were performed by using the following computing setup: a PC powered by an 8 core Intel Xeon E5440 (2.83GHz) with 16 GB of RAM.

The results are shown in Tables 5, 6, 7 where each column refers to a randomized-phase dataset generated with different MS *t, r* parameters.

Table 5 contains the switch error rates for the data after a phase randomization (mimicking the output of a genotyping process) which is used as a reference to evaluate the effectiveness of the phasing algorithms.

**Table 5.** Switch error (%) of the randomized sample: Each column refers to a randomized dataset generated with different MS *t, r* parameters whereas each value is the result of computing first the switch error in every window and then calculating their average

| (t, r) | 4, 0 | 4, 4 | 4, 40 | 40, 0 | 40, 4 | 40, 40 |
| --- | --- | --- | --- | --- | --- | --- |
| Switch Error | 11.24 | 10.36 | 9.54 | 9.83 | 11.24 | 10.55 |

**Table 6.** Phased switch error (%_0_): Each column refers to a dataset generated with different different mutation and recombination rate (MS *t, r* parameters). Each value is the average of the switch errors computed for every overlapping window.

| (t, r) | 4, 0 | 4, 4 | 4, 40 | 40, 0 | 40, 4 | 40, 40 |
| --- | --- | --- | --- | --- | --- | --- |
| Flash | 1.9 | 2.1 | 7.7 | 2.1 | 1.7 | 1.9 |
| Beagle | 2.6 | 1.4 | 2.0 | 0.7 | 0.6 | 0.6 |

**Table 7.** Processing time (s): Each column refers to a phased dataset. Each value in the Flash and Beagle rows is the time difference between the start of the application and the writing of the data to the storage. The speedup row contains the relative speedup of Flash with respect to Beagle.

| (t, r) | 4, 0 | 4, 4 | 4, 40 | 40, 0 | 40, 4 | 40, 40 |
| --- | --- | --- | --- | --- | --- | --- |
| Flash (s) | 0.05 | 0.05 | 0.05 | 0.24 | 0.25 | 0.24 |
| Beagle (s) | 4.28 | 2.23 | 2.24 | 11.05 | 11.64 | 13.17 |
| Speedup | 85.6 | 44.6 | 44.8 | 47.9 | 46.6 | 54.9 |

The randomized datasets were given to Beagle and Flash to reconstruct the phase information. Table 6 shows the switch error rate after the application of both phasing methods. Each value is the average of the switch errors computed for every overlapping window.

The Table 7 shows the computation time taken for each test and the relative speedup of Flash with respect to Beagle (last row).

#### 4.1.2. Real data

Real data are taken from the Genome 1000 project [1]. In particular we use phased data from the phase1, release v3 20101123 chromosome 20.

As described in the previous section, the Beagle tests were performed slicing the data before and after the computation. We named the first one *Beagle Whole* and the second *Beagle Sliced*.

The computing setup for Flash and Beagle Whole test was composed by a single node PC with an Intel Q6600 quadcore CPU (2.40GHz), 8GB RAM. The Beagle Sliced one was performed in a parallel environment using a Pydoop based application [9] as launcher. The parallel computing setup was composed by a cluster of 16 nodes with the same specification of the PC used for the simulated data test, for every node.

The Genome 1000 chromosome phased data contains information of different populations and obtained by different genotyping technologies. Therefore, to perform the haplotyping tests, we divided the chromosome data into the different populations and took the SNPs common to all the individuals. This data was then phase-randomized and used as input for the successive tests.

Flash runs were performed with two different configurations (the first one with a threshold of 100% both for solved individuals and computation effort whereas the second one with a threshold of 98% for the solved individuals and 10^−6^% for the computation effort) as explained in section 2.1.

The Beagle command line was the same for both Whole and Sliced runs (java -Xmx2048m with the remaining parameters left to the default value).

Table 8 shows the accuracy results. Each row refers to a different population. Flash Th and Flash NoTh columns show the Flash execution with and without threshold. The Beagle Wh column is related to the Beagle Whole run whereas the Beagle S1 refers to the sliced input dataset with each slice processed in parallel by a single Beagle istance.

**Table 8.** Beagle-Flash Switch error comparison: The mean switch error among the 5 SNP sized windows is shown. Flash NoTh and Flash Th results refer to the Flash execution with and without threshold. The Beagle Wh column is related to the Beagle Whole run whereas the Beagle SI refers to the sliced input dataset with each slice processed in parallel by a single Beagle instance.

| Dataset | Ind. | SNPs | Unphased SE (%) | FLASH NoTh SE (%) | FLASH Th SE (%) | Beagle Wh SE (%) | Beagle Sl SE (%) |
| --- | --- | --- | --- | --- | --- | --- | --- |
| ASW | 61 | 407230 | 14.55 | 3.10 | 2.82 | 1.80 | 3.52 |
| CEU | 85 | 243080 | 20.27 | 2.23 | 1.82 | 1.29 | 2.41 |
| CHB | 97 | 219825 | 20.87 | 2.14 | 1.71 | 1.27 | 2.11 |
| CHS | 100 | 219898 | 20.88 | 2.09 | 1.72 | 1.26 | 2.11 |
| CLM | 60 | 298412 | 17.69 | 3.47 | 2.59 | 2.14 | 3.38 |
| FIN | 93 | 234696 | 20.87 | 2.09 | 1.74 | 1.10 | 2.36 |
| GBR | 89 | 244783 | 20.19 | 2.27 | 1.83 | 1.26 | 2.39 |
| IBS | 14 | 172651 | 26.58 | 5.90 | 5.90 | 6.48 | 10.97 |
| JPT | 89 | 213243 | 21.42 | 2.11 | 1.77 | 1.28 | 2.23 |
| LWK | 97 | 438268 | 13.53 | 2.42 | 2.16 | 1.14 | 2.76 |
| MXL | 66 | 274042 | 18.66 | 3.21 | 2.33 | 1.83 | 3.03 |
| PUR | 55 | 306964 | 17.61 | 3.45 | 2.71 | 2.12 | 3.52 |
| TSI | 98 | 261117 | 19.03 | 2.14 | 1.71 | 1.23 | 2.22 |
| YRI | 88 | 398798 | 14.78 | 2.37 | 2.14 | 1.10 | 2.93 |

Table 9 shows the speed results. This test lacks the Beagle Sliced result because the computation of each window introduces a large overhead hindering any parallel execution advantage(i.e. actual results show that for a single window of 5 SNPs Beagle takes about 1 second on average leading roughly to 4000 seconds for the ASW dataset on a 100 nodes parallel run).

**Table 9.** Beagle-Flash performance comparison: Computation time. Flash NoTh and Flash Th results refer to the Flash execution with and without threshold. Beagle Whole is related to the Beagle execution with the whole dataset as input. The tow last columns show the speedup obtained with the two configurations of Flash with respect to Beagle.

| Dataset | Ind. | SNP's | Flash C/T NoTh(s) | Flash C/T Th(s) | Beagle Whole C/T (s) | Su NoTh | Su Th |
| --- | --- | --- | --- | --- | --- | --- | --- |
| ASW | 61 | 407230 | 47 | 26 | 2130 | 45.3 | 81.9 |
| CEU | 85 | 243080 | 45 | 21 | 1836 | 40.8 | 87.4 |
| CHB | 97 | 219825 | 44 | 21 | 1511 | 34.3 | 71.9 |
| CHS | 100 | 219898 | 46 | 29 | 1515 | 32.9 | 52.2 |
| CLM | 60 | 298412 | 50 | 35 | 1098 | 22.0 | 31.4 |
| FIN | 93 | 234696 | 59 | 25 | 1622 | 27.5 | 64.9 |
| GBR | 89 | 244783 | 48 | 21 | 1480 | 30.8 | 70.5 |
| IBS | 14 | 172651 | 111 | 109 | 107 | 1.0 | 1.0 |
| JPT | 89 | 213243 | 43 | 19 | 1246 | 29.0 | 65.6 |
| LWK | 97 | 438268 | 67 | 40 | 3635 | 52.2 | 90.9 |
| MXL | 66 | 274042 | 66 | 19 | 1144 | 17.3 | 60.2 |
| PUR | 55 | 306964 | 59 | 22 | 991 | 16.8 | 45.0 |
| TSI | 98 | 261117 | 51 | 25 | 1826 | 35.8 | 73.0 |
| YRI | 88 | 398798 | 52 | 33 | 2804 | 53.9 | 85.0 |

As stated in section 2.1 using Flash with the threshold makes the execution faster. The accuracy also improves. This is probably due to a constraint relax of the heuristic step of the algorithm when the threshold option is selected. Without threshold, the algorithm tries to solve all the individuals using the least number of haplotypes and, among the solutions with the same size, the most likely is selected. Analyzing the solutions in detail we found that this behavior forces the inclusion of haplotypes leading to an increased entropy (of the haplotype set) thus to a worse accuracy. Forcing a threshold focuses the haplotype selection on the most similar individuals thus lowering the entropy. The remaining individuals are solved using the most likely diplotype among the their compatible ones.

As expected *BeagleWhole* accuracy is by far better than *BeagleSliced* that shows lower accuracy also with respect to Flash.

Both simulated and real data show that Beagle is less accurate than Flash but slower (with the exception of the IBS dataset that is not so significant given the accuracy results for both the applications). With respect to different phasing uses (for instance the imputation of low resolution phased data), when dealing with association studies a lower accuracy could be less important. The speed advantage of Flash could be therefore a very attractive feature for GWAS.

### 4.2. Association test

To perform the association test the haplotypes for the whole population are selected by means of a sliding window approach into many overlapping stripes of fixed size.

The data set used is related to the chromosome 10 phenotyped for the type 1 diabetes. It is composed by 8286 SNPs and 5554 IDs divided in 3894 controls and 1660 cases.

To compare Flash with the well known Plink application, a standard *χ*^2^ test is performed on every window using a multimarker approach with 5 consecutive SNPs. Moreover we create different datasets, derived from the previous one by randomizing the phenotype information to create a worst case benchmark for the association. The comparison results are shown in figure 4.1. Flash, Plink and randomized phenotype results are depicted from the bottom to the top.

**Figure 4.1.**
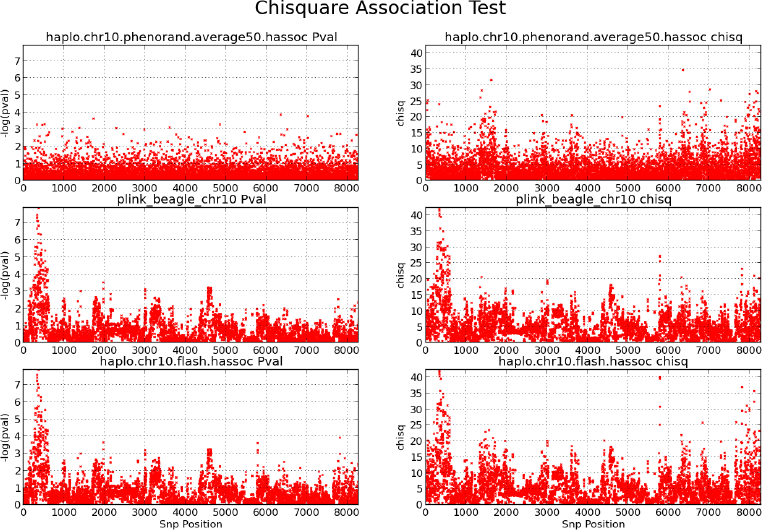
*χ*^2^ association test comparison among randomized phenotype (top), Plink (center), and Flash (bottom). Left pane shows the p-value. Right pane is for the *χ*^2^ value.

As expected, the association test performed on the randomized datasets does not produce any meaningful result, while Flash and Plink show a very similar behavior both finding a p-value of ∼ 10^−8^ in a region of a known association, but Flash is able to get these results with a speedup of about an order of magnitude.

To evaluate the power of a multimarker association test with respect to the single SNP one, we performed the 1-SNP test using Plink and compared the result with the Flash multimarker one for the same dataset. As can be seen in figure 4.2, the single SNP approach can resolve association in the same region with a smaller statistical power obtaining a p-value of about 10^−5^.

**Figure 4.2.**
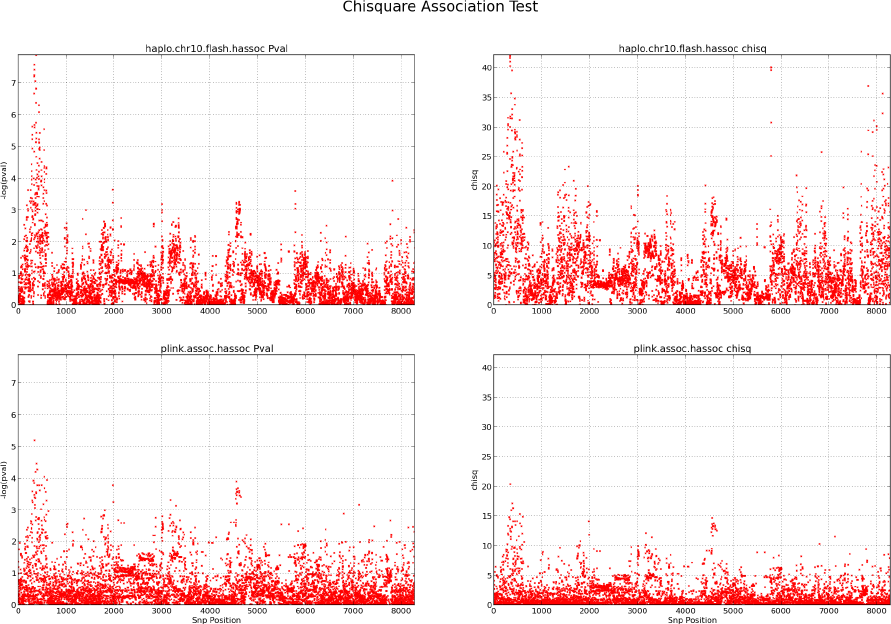
Flash (top) multimarker association test and single SNP with Plink (bottom) . Left pane shows the p-value. Right pane is for the *χ*^2^ value.

## 5. Conclusions

In this report we present Flash, an entropy minimization algorithm to infer local haplotypes of unrelated individuals. When compared with some state of the art applications such as BEAGLE[5] and MACH[10] it shows a speedup in execution time while maintaining a comparable accuracy. Finally we tested the statistical power of haplotype-based Genome Wide Association Study (GWAS) making a comparison with a single SNP association test performed with Plink [12]. Our tests show that the multimarker association test in this case is more powerful than the single SNP one as often reported in the literature. Moreover, Flash and Plink show similar results for the multimarker association test but Flash speeds up the computation time of about an order of magnitude using 5 SNP size haplotypes.

## Acknowledgements

We would like to thank Mario Falchi (Department of Genomics of Common Disease, School of Public Health, Imperial College London, Hammersmith Hospital, London, UK) for helpful suggestions, Serena Sanna (Istituto di Ricerca Genetica e Biomedica of the National Research Council, IRGB-CNR) and Ilenia Zara (Advanced Genomics Computing Technology group, CRS4) for useful discussions.

## Footnotes

1 http://ftp.1000genomes.ebi.ac.uk/voll/ftp/release/20110521/

## References

[1] Gongalo R Abecasis, David Altshuler, Adam Auton, Lisa D Brooks, Richard M Durbin, Richard a Gibbs, Matt E Hurles, and Gil a McVean. A map of human genome variation from population-scale sequencing. Nature, 467(7319):1061–73, October 2010.

[2] J Akey, L Jin and M Xiong. Haplotypes vs single marker linkage disequilibrium tests: what do we gain? European journal of human genetics: EJHG, 9(4):291–300, April 2001.

[3] David H Ballard, Judy Cho, and Hongyu Zhao. Comparisons of multi-marker association methods to detect association between a candidate region and disease. Genetic epidemiology, 34(3):201–12, April 2010.

[4] Paola Bonizzoni, Gianluca Vedova, Riccardo Dondi, and Jing Li. The Haplotyping problem: An overview of computational models and solutions. Journal of Computer Science and Technology, 18(6):675–688, November 2003.

[5] Sharon R Browning and Brian L Browning. Rapid and accurate haplotype phasing and missing-data inference for whole-genome association studies by use of localized haplotype clustering. American journal of human genetics, 81(5):1084–97, November 2007.

[6] Paul I W de Bakker, Roman Yelensky, Itsik Pe’er, Stacey B Gabriel, Mark J Daly, and David Altshuler. Efficiency and power in genetic association studies. Nature genetics, 37(11):1217–23, November 2005.

[7] Eran Halperin and Richard M. Karp The minimum-entropy set cover problem. Theoretical Computer Science, 348(2–3):240–250, December 2005.

[8] RR Hudson. Generating samples under a Wright-Fisher neutral model of genetic variation. Bioinformatics, 18(2):337–3382002.

[9] Simone Leo and Gianluigi Zanetti. Pydoop: a Python MapReduce and HDFS API for Hadoop. In Proceedings of the 19th ACM International Symposium on High Performance Distributed Computing, HPDC ‘10, pages 819–825, New York, NY, USA, 2010. ACM.

[10] Yun Li, Cristen J Willer, Jun Ding, Paul Scheet, and Gonçalo R Abecasis. MaCH: using sequence and genotype data to estimate haplotypes and unobserved genotypes. Genetic epidemiology, 34(8):816–34, December 2010.

[11] Aaron J Lorenz, Martha T Hamblin, and Jean-Luc Jannink. Performance of single nucleotide polymorphisms versus haplotypes for genome-wide association analysis in barley. PloS one, 5(ll):el4079, January 2010.

[12] Shaun Purcell, Benjamin Neale, Kathe Todd-Brown, Lori Thomas, Manuel a R Ferreira, David Bender, Julian Maller, Pamela Sklar, Paul I W de Bakker, Mark J Daly, and Pak C Sham. PLINK: a tool set for whole-genome association and population-based linkage analyses. American journal of human genetics, 81(3):559–75, September 2007.

[13] X. Wang, N. J. Morris, D. J. Schaid, and R. C. Elston. Power of Single-vs. Multi-Marker Tests of Association. Genet. Epidemiol, 5(36):480–487, January 2010.

